# Flexible adaptation of task-positive brain networks predicts efficiency of evidence accumulation

**DOI:** 10.1101/2023.09.07.556742

**Authors:** Alexander Weigard, Mike Angstadt, Aman Taxali, Andrew Heathcote, Mary M. Heitzeg, Chandra Sripada

## Abstract

Efficiency of evidence accumulation (EEA), an individual’s ability to selectively gather goal-relevant information to make adaptive choices, is thought to be a key neurocomputational mechanism associated with cognitive functioning and transdiagnostic risk for psychopathology. However, the neural basis of individual differences in EEA is poorly understood, especially regarding the role of largescale brain network dynamics. We leverage data from over 5,000 participants from the Human Connectome Project and Adolescent Brain Cognitive Development Study to demonstrate a strong association between EEA and flexible adaptation to cognitive demand in “task-positive” frontoparietal and dorsal attention networks, which explains 36%-39% of the variance across individuals in EEA. Notably, individuals with higher EEA displayed divergent task-positive network activation across n-back task conditions: higher activation under high cognitive demand (2-back) and lower activation under low demand (0-back). These findings suggest that brain networks’ flexible adaptation to cognitive demands is a key neural underpinning of EEA.

## Introduction

Evidence accumulation models^1^ posit that individuals complete many cognitive tasks by gradually accumulating noisy evidence for each possible choice until evidence for one choice reaches a critical threshold. This class of formal models has been highly successful at explaining key features of choice response time data and is now considered one of the predominant mathematical frameworks for modeling task performance across a wide variety of cognitive domains in the psychological and neural sciences^1–3^.

A growing literature has recently begun to reveal how the latent psychological mechanisms posited by evidence accumulation models contribute to higher-order cognition and behavior. Efficiency of evidence accumulation (EEA), or the ability to selectively accumulate goal-relevant evidence to make adaptive choices in the context of noisy information, appears to be a task-general process and a key underpinning of higher-order cognitive functions, including working memory and general intelligence^4–9^. In parallel, recent applications of these models in clinical research have demonstrated that EEA is significantly reduced across multiple disorders, including attention-deficit/hyperactivity disorder (ADHD)^10–13^, schizophrenia^14,15^, bipolar disorder^16^, and problematic substance use^17^, suggesting that lower EEA is a transdiagnostic cognitive risk factor for psychopathology^16,18^.

EEA’s relevance to cognitive functioning and psychiatric disorders has led to increasing interest in identifying its neural underpinnings. Experimental work in non-human primates has documented ramping patterns in neural firing rates during decision-making that display properties consistent with evidence accumulation processes^2,19–23^ and parallel signatures have been identified in humans using electroencephalogram (EEG)^24–29^ and functional magnetic resonance imaging (fMRI)^30–33^. Although multiple brain areas appear to be involved, converging evidence suggests that the frontoparietal network (FPN), a group of brain regions associated with task performance and cognitive control, plays a central role^34^. Outside of this experimental literature, recent studies using disparate methodologies have found that better EEA is associated with activation in the inferior parietal lobe during decision-making^35^, greater error-related activations in the salience network^36^, and a marker of neural speed derived from several EEG components^37^. Additionally, a recent multimodal neuroimaging investigation that focused on the dorsal portion of the FPN found evidence that white matter macrostructure within this subnetwork and its functional coupling with premotor cortex were both related to EEA^38^.

Despite the critical importance of this body of work for understanding the neural basis of EEA, a key limitation is that each of these studies had a constrained focus on specific brain regions or narrow subnetworks of regions. Recent research leveraging multivariate predictive modeling in neuroimaging data has demonstrated that many cognitive and psychological variables are only weakly associated with activity in discrete regions and are more robustly predicted by features of largescale brain networks that are distributed across the cortex^39–41^. Further, the properties of such brain networks are far from static, and instead display dynamic adaptations and reconfigurations to meet task demands^42,43^. Hence, although there is growing evidence that EEA may be a key neurocomputational underpinning of cognitive and adaptive functioning, its associations with the dynamic properties of largescale brain networks remain unclear.

In the current study, we present novel evidence that one such property shows a strong and robust association with EEA: the degree to which “task-positive” brain networks flexibly adapt to cognitive demand. The FPN and the dorsal attention network (DAN), another group of brain regions associated with the top-down control of attention^44^, are collectively labeled “task-positive” networks^45^ because they reliably show increased activity in task conditions that are cognitively demanding (i.e., difficult). As EEA is a formal measure of the ratio of task-relevant signal to task-irrelevant noise during cognitive processing^46,47^, it is conceptually linked to the interrelated functions of the FPN, which appears to selectively facilitate goal-relevant behaviors during task performance, and the DAN, which appears to modulate attentional resources toward goal-relevant information.

Parametric effects of cognitive demand on activity in the FPN and DAN are reliably observable during the commonly used n-back fMRI paradigm^48,49^, in which the difficulty of the cognitive task varies as a function of how many stimuli must be actively maintained in working memory to make accurate choices. Previous work has shown that higher levels of difficultly on the n-back generate neural activation maps that are more closely associated with cognitive abilities than those generated from less difficult n-back conditions^50^, suggesting that the degree to which individuals’ brain networks respond to the demands of a given task may have important implications for task performance. As flexible adaptation of neural systems to the demands of external tasks has long been theorized to support efficient cognitive processing^46,51^, we sought to directly assess the degree to which demand-related changes in neural activation across the FPN and DAN are associated with EEA.

Across two large data sets spanning different developmental periods, the Human Connectome Project (HCP)^52^ and the baseline sample of 9- and 10-year-old youth from Adolescent Brain Cognitive Development^SM^ Study (ABCD Study®)^53^, we first use multivariate predictive modeling to demonstrate that neural response to cognitive demand during the n-back explains a substantial portion (36%-39%) of the variance in individuals’ EEA on the task. We then show that this predictive relationship can be largely attributed to EEA’s association with demand-related activation patterns in the FPN and DAN. Critically, we provide novel evidence that this network configuration shows divergent relations with EEA under different levels of cognitive demand; although activation in task-positive networks during the difficult (2-back) condition is strongly positively related to EEA, activation in these networks during the easy (0-back) condition is strongly *negatively* related to EEA. These findings suggest that flexible adaptation to cognitive demands across task-positive brain networks is a key neural underpinning of EEA and its downstream consequences for cognition and behavior.

## Results

### 1. Neural responses to cognitive demand during the n-back explain a sizable proportion of the variance in individuals’ EEA on the task

We built multivariate models that used vertex-wise brain activation data from the n-back’s cognitive load (2-0) contrast to predict EEA metrics during the n-back task (see Methods for details on EEA metrics). We tested their generalizability in independent data using leave-one-site-out cross-validation^40^ in ABCD and 10-fold cross-validation in HCP. All analyses were adjusted for age, sex, race/ethnicity, and motion (framewise displacement) using the partial correlation technique described in Methods.

Neural responses to cognitive demand explained a large proportion of the variance across all measures of EEA in both samples (Figure 1A) and performance was consistently high across all ABCD sites and HCP cross-validation folds (Figure 1B). Performance of the models was highest when predicting the average of EEA across n-back load conditions, explaining 39% of the variance in ABCD and 36% of the variance in HCP. Predictions of EEA on the 0-back (ABCD = 32%, HCP = 35%) were slightly more accurate than predictions of EEA on the 2-back condition (ABCD = 30%, HCP = 26%).

**Figure 1.**
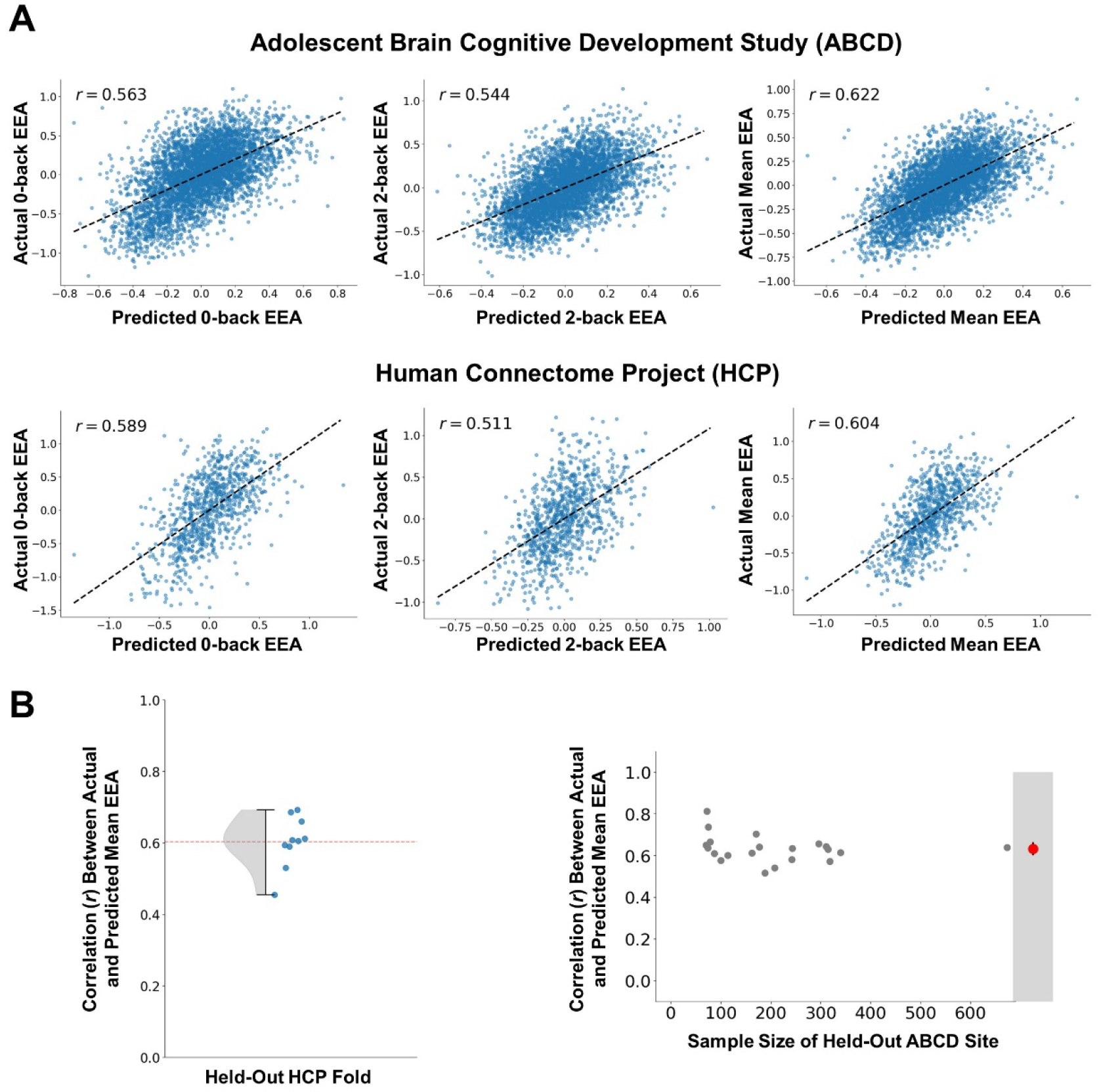
Relations between brain activation in the n-back cognitive load (2-0) contrast and efficiency of evidence accumulation (EEA). **A**) Correlations between EEA values predicted by the multivariate model and actual EEA values for the 0-back task, 2-back task, and the mean across both tasks in the Adolescent Brain Cognitive Development Study (ABCD) and Human Connectome Project (HCP) samples. The predicted values are drawn from models fit to independent data using the leave-one-site-out and 10-fold cross-validation methods in the ABCD and HCP samples, respectively. All values are residuals from regressions that adjusted for age, sex, race/ethnicity, and motion covariates. **B**) Correlations between predicted and actual mean EEA values in each of the 10 HCP test folds and each of the ABCD sites. For HCP, the density plot represents the distribution of values, and the red line represents the average value. For ABCD, the relation between correlations and sample size at each study site is displayed and the average value is displayed in the gray portion at right along with its 95% confidence interval.

This general pattern indicates that neural responses to cognitive demand are strongly related to measures of EEA across both levels of n-back load. Combined with the large observed correlations between EEA measured on the 0- and 2-back tasks (ABCD *r* = 0.45, CI = 0.42-0.48; HCP *r* = 0.54, CI = 0.48 – 0.59), these results are consistent with the hypothesis that EEA reflects a domain-general latent process that drives performance across tasks of both low and high complexity and has common neural underpinnings regardless of specific task demands^18^.

### 2. Features predictive of EEA show substantial overlap with the task-positive network regions activated in the n-back’s standard cognitive load contrast

Brain-wide consensus maps of feature weights from the models predicting EEA with activation in the cognitive load (2-0) contrast showed a strong visual similarity to the same contrast’s group-average activation maps (Figure 2). As expected, regions in the FPN and DAN were heavily represented across both types of maps. Most of the prefrontal and midline regions strongly activated by the load contrast were also heavily featured in the predictive model, although there were some apparent differences between the maps in lateral parietal regions. These spatial patterns were remarkably consistent across the ABCD and HCP samples. However, one notable difference between the samples is the finding of generally lower effect sizes in the group-average 2-0 activation map in ABCD relative to HCP, which could indicate that while children and adults activate similar networks during high cognitive demand, activation levels are generally lower in children compared to adults, consistent with recent findings^54^.

**Figure 2.**
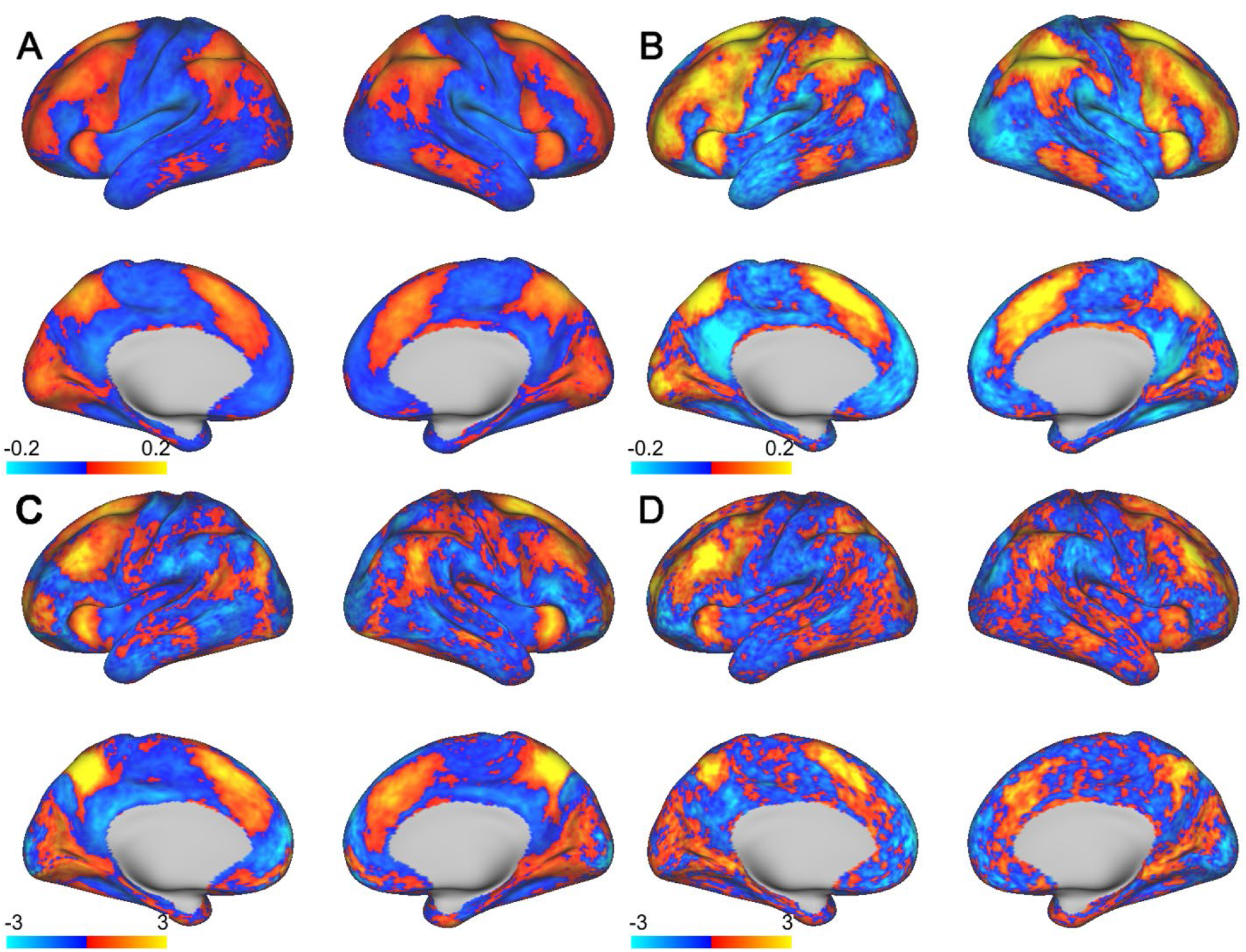
Group-level cortical maps of effect sizes (Cohen’s d) from the n-back cognitive load (2-0) contrast and feature weights (converted to Z-scores: mean = 0, SD = 1) from the models predicting individuals’ n-back task performance with this contrast. **A**) Effect size map for the 2-0 contrast in the Adolescent Brain Cognitive Development Study (ABCD) sample. **B**) Effect size map for the 2-0 contrast in the Human Connectome Project (HCP) sample. **C**) Consensus feature weight map for models predicting performance in the ABCD sample. **D**) Consensus feature weight map for models predicting performance in the HCP sample.

### 3. Task-positive network activations during the 2-back and 0-back make significant contributions to prediction of EEA

We sought to further parse out the role of specific task-positive networks, and their dynamic changes across levels of cognitive demand, in predicting EEA. We therefore examined associations among EEA, 0-back and 2-back activations in the DAN and FPN (averaged across the entire networks’ Gordon parcellations), and load-related differences in DAN and FPN activations (2-back minus 0-back) for both ABCD and HCP (Table 1). Load-related differences (2-0) in the FPN and DAN showed the expected positive relations with EEA, consistent with the idea that neural responses to cognitive demand in both the FPN and DAN make key contributions to the fMRI data’s predictive associations.

**Table 1.** Adolescent Brain Cognitive Development Study (ABCD) and Human Connectome Project (HCP) correlations between efficiency of evidence accumulation (EEA) and average measures of frontoparietal network (FPN) and dorsal attention network (DAN) activation for the 0-back, 2-back and cognitive load (2-0) contrast. All variables were adjusted for age, sex, race/ethnicity, and motion (framewise displacement) by using multiple regression to remove variance associated with these covariates. 95% confidence intervals, displayed in italics next to each correlation, were estimated using a clustered bootstrapping procedure that accounted for nesting by family and study site.

|  | ABCD<br>*r* | ABCD<br>95% CI | HCP<br>*r* | HCP<br>95% CI |
| --- | --- | --- | --- | --- |
| FPN 0 | -.27 | -.24 -.29 | -.31 | -.24 -.37 |
| FPN 2 | .17 | .22 .13 | .08 | .15 .02 |
| FPN 2-0 | .36 | .39 .33 | .46 | .51 .41 |
| DAN 0 | -.07 | -.04 -.09 | -.14 | -.07 -.21 |
| DAN 2 | .17 | .21 .14 | .08 | .15 .00 |
| DAN 2-0 | .22 | .24 .19 | .30 | .35 .23 |

### 4. Task-positive network activations during the 2-back and 0-back are positively correlated with one another but show strongly divergent relations with EEA

For both task-positive networks, there were strong positive correlations between each subject’s activation in the 2-back condition and their activation in the 0-back condition, both in the HCP sample (FPN *r* = 0.64, CI = 0.60 – 0.68; DAN *r* = 0.74, CI = 0.71 – 0.77) and in the ABCD sample (FPN *r* = 0.24, CI = 0.21 – 0.27; DAN *r* = 0.36, CI = 0.33 – 0.38). These strong dependencies were notable given that we observed strongly *divergent* relationships between activation in the 2-back and 0-back conditions and EEA (Table 1). More specifically, 2-back task-positive network activation is positively related to EEA while 0-back activation is negatively related to EEA. Figure 3 illustrates these complex interrelations by plotting each individual’s 0-back activation levels on the x-axis against their 2-back activation levels on the y-axis. The strong, positive relationship between 2-back and 0-back activation is shown by the black dashed regression line. For both ABCD and HCP, and for both task-positive networks (FPN and DAN), we observed a common pattern: individuals in the upper left-hand quadrant, who have relatively greater activation in the 2-back and lower activation in the 0-back condition (i.e., relative to the regression line) show the highest EEA, which is indicated both by the darker red hue of the points as well as the mean standardized EEA scores displayed in the quadrant. Individuals in the lower right-hand quadrant, who have relatively lower 2-back activation and higher activation in the 0-back condition (relative to the regression line), show the lowest EEA.

**Figure 3.**
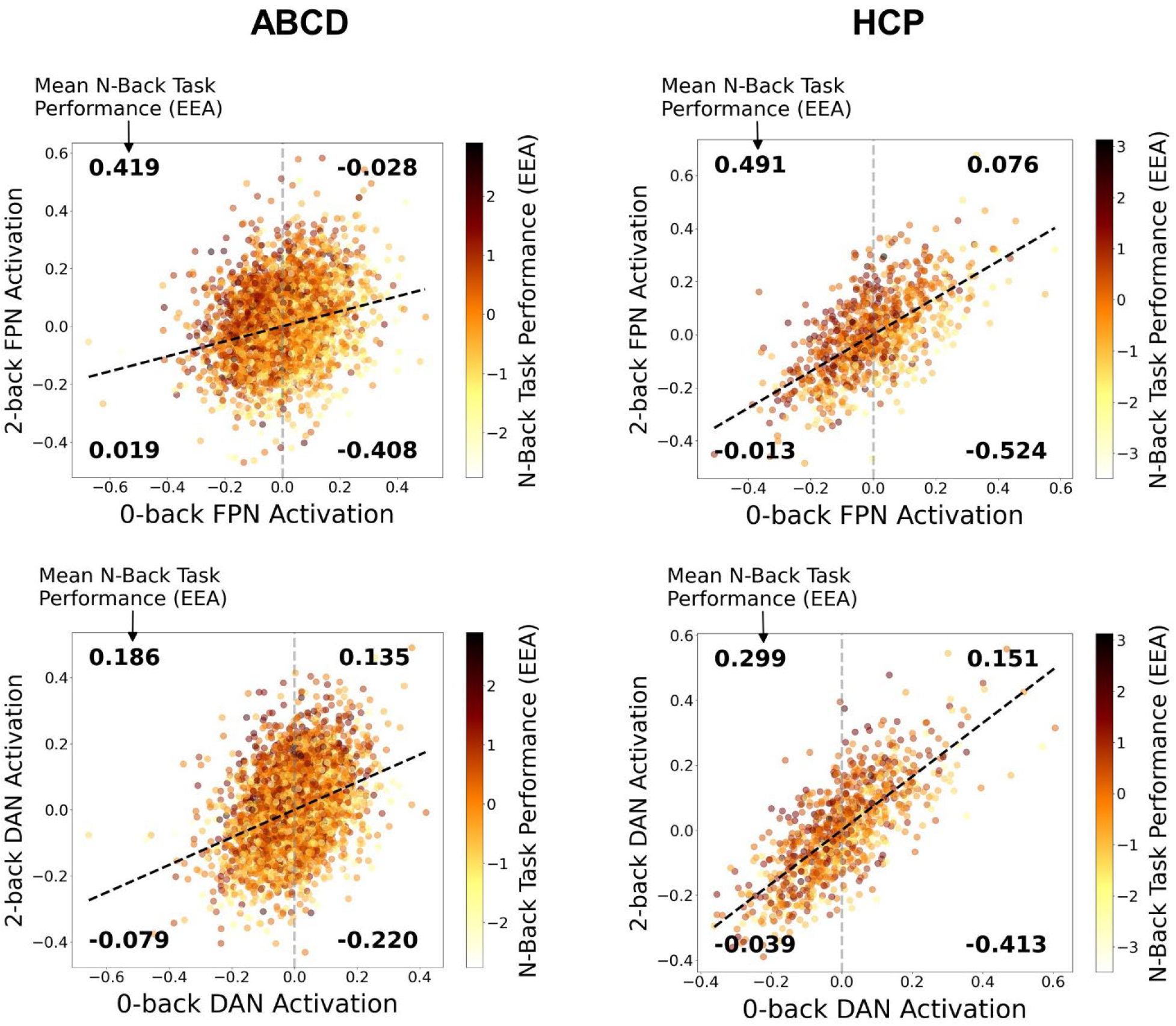
Visualization of dynamic relations between average task-positive network activation in the 0-back and 2-back conditions and overall efficiency of evidence accumulation (EEA) on the task for the Adolescent Brain Cognitive Development Study (ABCD; left column) and Human Connectome Project (HCP; right column) samples. All values were adjusted for age, sex, race/ethnicity, and motion covariates and were then converted to standardized scores (Z-scores: mean = 0, SD = 1) for interpretability. Individuals’ EEA is represented by the hue of the points, with individuals higher in EEA having darker red hues. Activations of the frontoparietal network (FPN) are shown in the top row while activations of the dorsal attention network (DAN) are shown in the bottom row. Black dotted lines represent the regression line for relations between 0-back and 2-back task activations. Combined with the gray dotted lines representing the average 0-back activation level, the regression lines form four quadrants that denote whether individuals have higher or lower 2-back activation than would be expected given their level of 0-back activation. Bold numbers reflect the average EEA of individuals in each quadrant.

The preceding results highlight that, because of the positive dependency between activations across different levels of demand, the absolute level of FPN and DAN activation during a given task condition is relatively unimportant for predicting EEA. Rather, the more important factor is the *differences* in activation between high demand and low demand conditions. Individuals for whom this difference is large simultaneously exhibit higher than expected activation in the high-demand (2-back) condition and lower than expected activation in the low-demand (0-back) condition, which appears to be the key signature of better EEA. Consistent with this idea, the correlation values reported in Table 1 indicate that variance in EEA explained by the 0-back and 2-back activations measured within individual load conditions is consistently lower than the variance explained by differences in activation between the high demand and low demand conditions (within condition: ABCD FPN = 10%; ABCD DAN = 3%; HCP FPN = 10%; HCP DAN = 3%; difference between conditions: ABCD FPN = 13%; ABCD DAN = 5%; HCP FPN = 21%; HCP DAN = 9%).

### 5. EEA is associated with flexibility in the engagement of the FPN and DAN across different levels of cognitive demand

The findings detailed above suggest that individuals with higher EEA on the n-back tend to be those whose task-positive networks show higher activity in high-demand conditions but lower activity in low-demand conditions. To directly investigate this possibility, we sorted individuals in each sample into 10 bins ordered by EEA and plotted each bin’s mean FPN and DAN activations (Figure 4). We also display regression lines for the association between bin order and activation to illustrate the overall pattern. Across both task-positive networks and both samples, greater EEA was associated with increases in network activity during the 2-back and corresponding decreases in network activity during the 0-back. Put another way, individuals with the lowest EEA engage these networks to a similar degree regardless of cognitive demands. In contrast, individuals with the highest EEA display the greatest degree of flexible modulation of task-positive network engagement in response to demands, exhibiting *both* the highest activation in task positive networks in the 2-back and the lowest activation in the 0-back.

**Figure 4.**
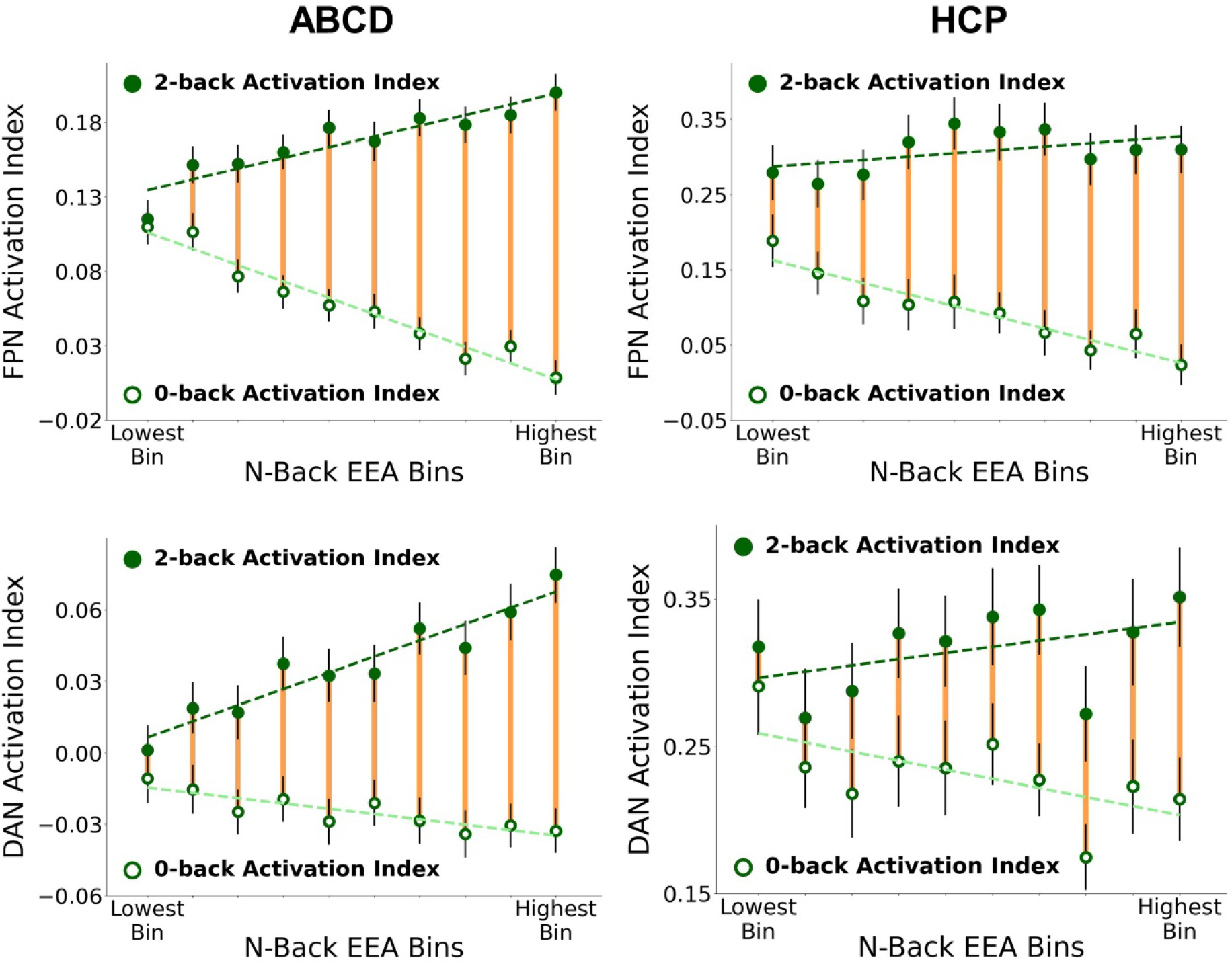
Plots of mean frontoparietal network (FPN; top row) and mean dorsal attention network (DAN; bottom row) activations in the 0-back and 2-back conditions for 10 bins of individuals ranked by their mean EEA in the Adolescent Brain Cognitive Development Study (ABCD; left column) and Human Connectome Project (HCP; right column) samples. Vertical green lines around each point represent 95% confidence intervals while yellow lines highlight the difference between 0-back and 2-back means. Dashed lines represent regression lines for the relation between bin rank and each brain activation index.

### 6. Differences between samples suggest potential developmental changes in task-positive network engagement

Although most brain network activation patterns and their associations with EEA were remarkably consistent across the child ABCD sample and adult HCP sample, several key differences alco emerged. As noted above, adults in HCP showed larger activation effects in the 2-0 contrast (despite showing similar spatial patterns of activation) than children in the ABCD sample. Adults also exhibited a much stronger positive dependency in task-positive network activations across high- and low-load conditions compared to children. In addition, the values displayed in Figure 4 suggest possible developmental differences in overall levels of DAN activation under low cognitive demands. DAN activations in the 2-back were generally positive on average for both samples. However, ABCD participants’ 0-back DAN activations were negligible or negative on average whereas HCP participants’ DAN activations were consistently positive on average. Although the methodological differences between the tasks and sampling methods in ABCD and HCP (Methods) preclude strong conclusions about developmental change, this pattern of results provides preliminary evidence that children do not consistently engage the DAN on low-load conditions and that improvements in performance in adulthood could be partially attributable to greater DAN engagement.

## Discussion

A growing literature on computational evidence accumulation models suggests that EEA, the rate at which individuals gather goal-relevant evidence to make adaptive choices, is a foundational mechanism that drives individual differences across many cognitive functions and has clear relevance to psychiatric disorders^5,6,18^. Although evidence accumulation processes are well-characterized at the level of discrete neurophysiological recordings during decision making^2,19–23,55^, the role of largescale brain networks in supporting individual differences in EEA remains poorly understood. The current study is the first to document a pattern of largescale brain network dynamics that shows strong and generalizable associations with EEA across two large, diverse samples of children and adults. We demonstrate that neural responses to cognitive demand on the n-back can account for a large portion of the variance in EEA on the task and that this association is largely driven by neural responses across the FPN and DAN, two “task-positive” networks involved in the control of attention and goal-directed cognition. Crucially, we find a divergent pattern in which individuals with higher EEA exhibit *both* higher activity in task-positive networks during a difficult task condition with high cognitive demands (2-back) as well as lower activity in these same networks during a less demanding task condition (0-back). Although these findings are consistent with prior work suggesting that evidence accumulation processes are supported by FPN regions^34,38^, the current study goes beyond this work in demonstrating a critical role for flexible adaptation of task-positive networks. That is, we demonstrated that dynamic changes in the activity of task-positive networks across different levels of cognitive demand, rather than these networks’ static properties, are closely linked to EEA. Indeed, FPN and DAN activation measures drawn from only single levels of cognitive demand showed systematically weaker associations with EEA than measures of differences in FPN and DAN activation across high- and low-demand levels.

This set of findings naturally raises the question of how these complex network dynamics relate to the process interpretation of EEA in the cognitive modeling framework. As EEA is a formal index of the extent to which individuals can selectively parse goal-relevant evidence from noise in order to make adaptive choices during behavioral tasks^18^, its opposing relations with task-positive network activity across the 0-back and 2-back conditions could reflect the modulation of attention. Specifically, during a difficult task that requires significant attentional resources (2-back), individuals with high EEA may allocate more attention to task-relevant features, resulting in greater 2-back activation in networks linked to attentional control and external task engagement. However, when the task at hand is relatively easy and does not require significant attentional resources to perform, individuals with high EEA may re-allocate attention away from the task at hand in order to preserve these resources, leading to decreased activation in the same networks. These complementary processes, the allocation of attention to tasks that strongly require it and the efficient redistribution of attention during those do not, both arguably reflect flexible modulation of attention in response to environmental demands.

Such an explanation is notably consistent with the established conceptualization of attentional modulation posited in “adaptive gain” theory^51^. This theory posits that the norepinephrine system supports cognitive performance by optimizing an individuals’ trade-off between exploitation of specific tasks and exploration of the larger environment. When task performance is not optimal, or in other words when an individual is failing to extract the maximum amount of reward from a task, adaptive gain theory posits that the norepinephrine system works in coordination with prefrontal cortical areas to increase neural signal-to-noise ratios through the allocation of attention to the task-relevant stimuli. However, when further allocation of attention to a task is less likely to increase rewards (e.g., on a less demanding task), the same systems serve to disengage attention from the task at hand in order to re-allocate cognitive resources to the exploration of the environment in search of more rewarding behaviors. As EEA in cognitive models is a formal representation of the signal-to-noise ratio in the decision process, adaptive gain theory has been invoked to explain findings of poorer EEA in ADHD as reflecting neural systems’ failure to flexibly modulate attention and arousal in response to external task demands^46,47^. The pattern of largescale brain network activation linked to EEA in the current study is consistent with this explanation, as it indicates that individuals lower in EEA engage networks involved in control of attention and external task processing to a similar degree regardless of task difficulty, whereas those with higher EEA flexibly modulate the engagement of these networks in response to changes in difficulty. Therefore, future work guided by adaptive gain theory may show promise for linking these network dynamics to the functioning of the norepinephrine system, especially given recent experimental findings indicating that norepinephrine agonists enhance individuals’ EEA^13,29,56^.

Our findings also bear interesting connections with previous theories regarding neural correlates of general cognitive ability. One notable connection is with a large literature that finds individual differences in general cognitive ability are linked to conflicting patterns of neural recruitment across several neuroimaging modalities, a pattern of findings that was reconciled in a review proposing that these effects are moderated by cognitive demand.^57^. More specifically, individuals with higher general cognitive ability show less recruitment of neural resources in lower-demand tasks (greater “neural efficiency”) but are also thought to recruit more resources in tasks with high demands^57^. As EEA appears to be a key driver of general cognitive ability^5,6^, the opposing relations between network activations at different levels of cognitive demand and EEA that are reported in the current study are broadly consistent with this theory. Importantly, this theory differs somewhat from the “adaptive gain” explanation detailed above; it posits that individuals with better cognitive performance differ in two distinct processes – greater neural efficiency on easy tasks and a separate tendency to allocate more neural resources to difficult tasks – rather than in the single process of flexible attention modulation. The current data cannot easily disambiguate between explanations, and future work would be necessary to do so. However, it should be noted that these explanations are not mutually exclusive and that the two-process explanation continues to imply greater flexibility in network modulation for individuals with higher EEA, as it posits that they both require fewer neural resources to perform a given task well and are also able to flexibly allocate more of these resources when more are needed.

Our results also highlight another somewhat surprising phenomenon: despite showing strongly opposing relations with EEA, task-positive activations in the high and low demand conditions are themselves positively correlated with one another, especially for adults in the HCP sample. This positive correlation may reflect a “general activation” dimension that represents individual differences in task-positive networks’ activation during most external tasks, regardless of these tasks’ level of demands. A key consequence of this finding is that activation in task-positive networks within single task conditions are more weakly related to EEA compared to differences in activation across high- and low-demand conditions. Therefore, from a practical standpoint of maximizing prediction of EEA, and perhaps other cognitive variables, from task-related neuroimaging data, it follows that optimal prediction may require explicit experimental manipulations of cognitive demands in order to measure network adaptation, which the current study suggests is the most robust predictor of EEA.

Although we focused on canonical networks derived from resting state connectivity, it is worth pointing out that our results are relevant to the growing body of work on the “multiple demand network” (MDN), a set of structures that shows considerable overlap with the FPN and DAN and that appears to support cognitive performance across a wide array of tasks^58,59^. The generality of this network’s relation to task behavior has led to the suggestion that the MDN is the basis of the common factor that explains covariance across many “executive function” tasks^54^. As EEA shows similar task-general properties and was recently demonstrated to be a strong explanation for the common factor of executive functioning^60^, future work on the MDN may be able to determine whether EEA mediates the relation between demand-related MDN dynamics and performance on diverse executive tasks. Our findings are also notably consistent with recent work demonstrating that children and adults tend to activate similar brain structures in the MDN during task performance but that children do so to a lesser degree^54^.

In conclusion, the current study characterizes dynamic properties of large-scale brain networks that show strong and replicable associations with EEA, a foundational cognitive individual dimension derived from a well-developed cognitive modeling literature. The findings specifically highlight individuals’ ability to flexibly engage, versus disengage, the FPN and DAN in conditions of high, versus low, cognitive demand as a key underpinning of EEA. These findings suggest several productive future avenues of investigation into relations among largescale brain network dynamics, neurotransmitter systems thought to support flexible behavior, and EEA’s downstream consequences for adaptive functioning and risk for clinical disorders.

## Methods

### HCP Sample and Data Acquisition

Data for the HCP sample was taken from the HCP-1200 release ^52,61^. Subjects provided informed consent, and all study procedures were approved by the Washington University Institutional Review Board.

Subjects completed two runs of an N-back working memory task (approximately 5 minutes each, TR=720ms, 2.0mm isotropic voxels). High resolution (0.7 mm isotropic) T1-weighted and T2-weighted images were also collected and used for data processing. Comprehensive details are available elsewhere on HCP’s overall neuroimaging approach ^52,62^ and HCP’s task fMRI dataset^63^.

Following exclusions for neuroimaging and behavioral data quality (described below), a total of 883 participants (465 females; mean age = 28.6, SD age = 3.7) were included in analyses.

### ABCD Sample and Data Acquisition

The ABCD Study® is a multisite longitudinal study with 11,875 children between 9 and 10 years of age from 22 sites across the United States^53^. The study conforms to the rules and procedures of each site’s Institutional Review Board, and all participants provide informed consent (parents) or assent (children). Data for this study are from ABCD Release 4.0.

High spatial (2.4 mm isotropic) and temporal resolution (TR=800 ms) resting state fMRI was acquired for the emotional N-back task in two separate runs (approximately 5 minutes each). High resolution (1 mm isotropic) T1-weighted and T2-weighted images were also collected and used for data processing.

Following exclusions for neuroimaging and behavioral data quality (described below), a total of 4,315 participants (2,182 females; mean age = 10.0, SD age = 0.63) were included in analyses.

### Neuroimaging Data Processing

Preprocessing was performed using fMRIPrep version 1.5.0^64^. Briefly, T1-weighted (T1w) and T2-weighted images were run through recon-all using FreeSurfer v6.0.1. Functional data were corrected for fieldmap distortions, rigidly coregistered to the T1, motion corrected, normalized to standard space, and transformed to CIFTI space with 91,282 grayordinates. All preprocessed data were visually inspected separately for the quality of the registration to the T1w image and the quality of the normalization to MNI space.

Task models were constructed following HCP scripts. Models were constructed using FSL (6.0.5.2) in a two-stage procedure, estimating each run, and then averaging runs. CIFTI data were smoothed with a 2mm FWHM Gaussian using HCP Connectome Workbench (1.4.2). Images were then high pass filtered at 0.005 Hz. Nuisance covariates in the first level models consisted of 24 motion correction parameters (3 rotation, 3 translation, first derivatives of each, and quadratics of original and derivatives), top 5 principal components of signal from white matter, top 5 principal components of signal from cerebrospinal fluid, and individual regressors for each TR that exceeded a 0.9mm framewise displacement. Task conditions modeled included 0-back and 2-back for each category of stimuli (HCP: faces, places, body parts, tools; ABCD: happy faces, neutral faces, fearful faces, places). Linear contrasts were constructed for contrasts of interest: 0-back, 2-back, 2-back vs 0-back.

Individual runs were considered good if they passed visual inspection and had at least 4 minutes of uncensored data. Subjects were only included if they had two good runs and complete task behavioral data that met the inclusion criteria described in the section below.

### EEA Estimation

EEA was estimated by fitting the diffusion decision model (DDM), a widely used evidence accumulation model^65^, to data from the n-back task in both the ABCD and HCP samples using Bayesian estimation methods implemented within the Dynamic Models of Choice (DMC)^66^ suite of R functions. In both samples, participants completed 80 trials in each of the two cognitive load conditions (0-back, 2-back). Specific details of the stimuli and task parameters are described in detail elsewhere^52,53,63^. At both levels of load, trials could be 1) “target” stimuli, which meet specific criteria for being targets (e.g., in the 2-back, stimuli that were previously presented exactly 2 spaces back), 2) “novel” stimuli, which are stimuli that have never been presented before, 3) “lure” stimuli, which were recently presented stimuli that do not meet the specific criteria for being targets. Lures are more difficult for participants to reject and ensure that participants are applying the full target criteria while completing the task rather than relying on the familiarity of stimuli, alone. The DDM included eight parameters for each level of cognitive load (0-back/2-back): three separate drift rate (*v*) parameters for target, novel, and lure stimuli, single boundary separation (*a*), non-decision time (*t0*), non-decision time variability (*st0*), and start point (*z*) parameters, and a parameter for the probability of “contaminant omissions”, which are non-responses due to causes outside of the main DDM response process (e.g., inattention)^67^. Omissions due to the task design (i.e., response cut off by the 2-second response window) were also addressed using methods developed in prior work on addressing omissions with evidence accumulation models^67^. The two other variability parameters included in the “full” DDM (*sv, sz*) were not estimated due to difficulties with accurately recovering these parameters at low numbers of trials.

Prior to estimation, we excluded individuals’ data if they displayed accuracy rates close to chance (<55%) or excessive rates of omissions/non-responses (>25%) in a given load condition, both of which indicate likely disengagement from the task. We also excluded RTs <200ms as these RTs are likely to reflect fast guesses by participants. Informative priors for parameter estimates were generated following a procedure we previously developed^68^. A hierarchical version of the DDM was fit to an independent sample of 300 ABCD participants who had failed neuroimaging data quality checks but not behavioral data quality checks and who were unrelated to the ABCD participants included in the main analyses of this study. Following parameter estimation for this independent subsample, we fit truncated normal distributions to the full distribution of all individual-level posterior samples for each parameter. These truncated normal distributions were then used as informative priors for model fits in the ABCD sample. For the HCP sample, we multiplied the scale of these priors by 1.5 to make them slightly less informative given that the adults in HCP likely display some developmental differences relative to the prior-generation sample drawn from children in ABCD.

The DDM was then estimated under informative priors at the individual level using the automated RUN.dmc() function that repeats the posterior sampling process until convergence is obtained (rhat <1.1). Convergence was corroborated by visually inspecting a subset of individuals’ sampling chains. Model fit was assessed using posterior predictive plots^69^, which indicated that the model provided an adequate description of the behavioral data in both samples (Supplemental Figures 1-2). Posterior medians for the drift rate (*v*) parameter were averaged across the three types of stimuli (target, novel, lure) to index individuals’ EEA at each level of cognitive load.

### Multivariate Predictive Modeling

Subject level contrast images for 2-back vs 0-back were used in a cross-validated principal components regression (PCR) predictive model^40^. In brief, this method performs dimensionality reduction on input data, fits a regression model on the resulting components, and applies this model out of sample in a 10-fold (HCP) or leave-one-site-out (ABCD) cross-validation framework. Nuisance covariates (age, age squared, sex, race/ethnicity, framewise displacement estimate of motion, framewise displacement squared) are handled by calculating a cross-validated form of partial correlation. In each training fold after the PCA is conducted to reduce the data, K components are retained, with the optimal value for K being estimated with a nested 5-fold cross-validation within just the training data. Both the component expressions as well as the outcome variable are regressed against nuisance variables. The betas estimated from this model are used to residualize both the training and test data, then a linear model is fit on the training data to predict the residualized outcome with the residualized expressions. This model is then applied to the test data to get a predicted value, which can then be correlated with the residualized outcome to get an out-of-sample partial correlation estimate. This is repeated for each fold, and then the per-fold correlations are averaged across folds.

### Network Activation Extractions

Average FPN and DAN activation values are calculated from the 0-back and 2-back contrast images separately. The contrasts are averaged within each network, using the 7-network parcellation from Yeo^70^.

### Relations Among Network Metrics and EEA

Following predictive modeling analyses and extractions of FPN and DAN network means, we assessed correlation coefficients (*r*) for relations among EEA and average 0-back, 2-back, and cognitive load contrast (2-back minus 0-back) activation values for each network. For all analyses involving EEA and these network summary variables, nuisance covariates (age, age squared, sex, race/ethnicity, framewise displacement estimate of motion, framewise displacement squared) were addressed by fitting multiple regression models in which each variable of interest was predicted by all nuisance covariates. Residuals from these models were used in all analyses and plots except for the binned EEA plots in Figure 4, which used raw values of all variables for interpretability. Sensitivity analyses revealed no substantive differences between inferences drawn from raw versus covariate-residualized values (Supplemental Materials; Supplemental Table 1; Supplemental Figure 3). A clustered bootstrapping procedure was used to estimate 95% confidence intervals (CIs) for *r* while accounting for nesting of individuals within families and ABCD sites.

## Acknowledgements

AW was supported by K23 DA051561. CS was supported by and the Dana Foundation David Mahoney Neuroimaging Program. AW, AH, and CS were supported by R01 MH130348. CS and MH were supported by U01 DA041106. AH was supported by Vidi grant VI.Vidi.191.091 from the Dutch Research Council.

Data used in the preparation of this article were obtained from the Adolescent Brain Cognitive Development^SM^ (ABCD) Study (https://abcdstudy.org), held in the NIMH Data Archive (NDA). This is a multisite, longitudinal study designed to recruit more than 10,000 children aged 9-10 and follow them over 10 years into early adulthood. The ABCD Study® is supported by the National Institutes of Health and additional federal partners under award numbers U01DA041048, U01DA050989, U01DA051016, U01DA041022, U01DA051018, U01DA051037, U01DA050987, U01DA041174, U01DA041106, U01DA041117, U01DA041028, U01DA041134, U01DA050988, U01DA051039, U01DA041156, U01DA041025, U01DA041120, U01DA051038, U01DA041148, U01DA041093, U01DA041089, U24DA041123, U24DA041147. A full list of supporters is available at https://abcdstudy.org/federal-partners.html. A listing of participating sites and a complete listing of the study investigators can be found at https://abcdstudy.org/consortium_members/. ABCD consortium investigators designed and implemented the study and/or provided data but did not necessarily participate in analysis or writing of this report. This manuscript reflects the views of the authors and may not reflect the opinions or views of the NIH or ABCD consortium investigators. The ABCD data repository grows and changes over time. The ABCD data used in this report came from ABCD release 4.0 (https://nda.nih.gov; DOI 10.15154/1,523,041).

## Data and Code Availability

The ABCD data used in this report came from ABCD release 4.0 (https://nda.nih.gov; DOI 10.15154/1,523,041). ABCD data specific to the current study can be accessed at NDA Study 2297 (DOI 10.15154/wnt8-dq37). HCP data used in this manuscript are accessible at https://github.com/SripadaLab. Code for all study analyses is available at https://osf.io/yte76/.

## Author Contribution Statement

Conceptualization: AW, CS, AT, MA, AH, MH; Methodology: AW, CS, AT, MA, AH; Formal analysis: AW, AT, MA; Data curation: MA, AT; Writing—original draft: AW; Writing—reviewing and editing: AW, CS, AT, MA, AH, MH; Visualization: AT, MA; Funding acquisition: MH, CS, AW.

