## Supplemental Materials for "Flexible adaptation of task-positive brain networks predicts efficiency of evidence accumulation"

**Diffusion Decision Model Fit**

Posterior predictive plots of the diffusion decision model (DDM) description of the Human Connectome Project (HCP) and Adolescent Brain Cognitive Development Study (ABCD) n-back task data are displayed in Supplemental Figures 1-2 below. The DDM provided an excellent description of accuracy rates across all conditions and generally provided an accurate description of response time latency, although it slightly overestimated several longer response time quantiles for target trials. Overall, the DDM provides an adequate description of trends in the choice response time data in both load conditions and in both samples.

**Sensitivity Analyses Without Covariate Correction**

We conducted sensitivity analyses for all primary analyses involving network mean measures without adjusting values for covariates in order to gauge the potential impact of our covariate controls on substantive inferences. Comparison of values in Supplemental Table 1 from these unadjusted analyses with those in Table 1 of the main manuscript indicates that, although most relationships between network mean measures and efficiency of evidence accumulation (EEA) appear to be slightly stronger without covariate adjustment, the general pattern of these relationships is nearly identical. Correlations between 0-back and 2-back activations in task-positive networks without covariate adjustment were practical identical to those reported in the main paper for both HCP (FPN *r* = 0.64, CI = 0.60 – 0.68; DAN *r* = 0.74, CI = 0.70 – 0.76) and ABCD (FPN *r* = 0.24, CI = 0.21 – 0.27; DAN *r* = 0.35, CI = 0.33 – 0.38). The dynamic relations between 0-back activation, 2-back activation and EEA also remained the same (Supplemental Figure 3). Taken together, these sensitivity analyses indicate that covariate corrections used in the main manuscript had little impact on the main substantive findings of the study.

**Supplemental Figure 1.** Posterior predictive plots for diffusion decision model (DDM) fits to the Human Connectome Project (HCP) n-back task data in each cognitive load and trial condition. Plots display the cumulative probability of a “non-target” (solid line) and “target” (dotted line) response for empirical (thick line) and model-predicted (thin line) data. Specific response time (RT) quantiles (.1, .3, .5, .7, .9) are also displayed for empirical (open dots) and model-predicted (solid dots) data.

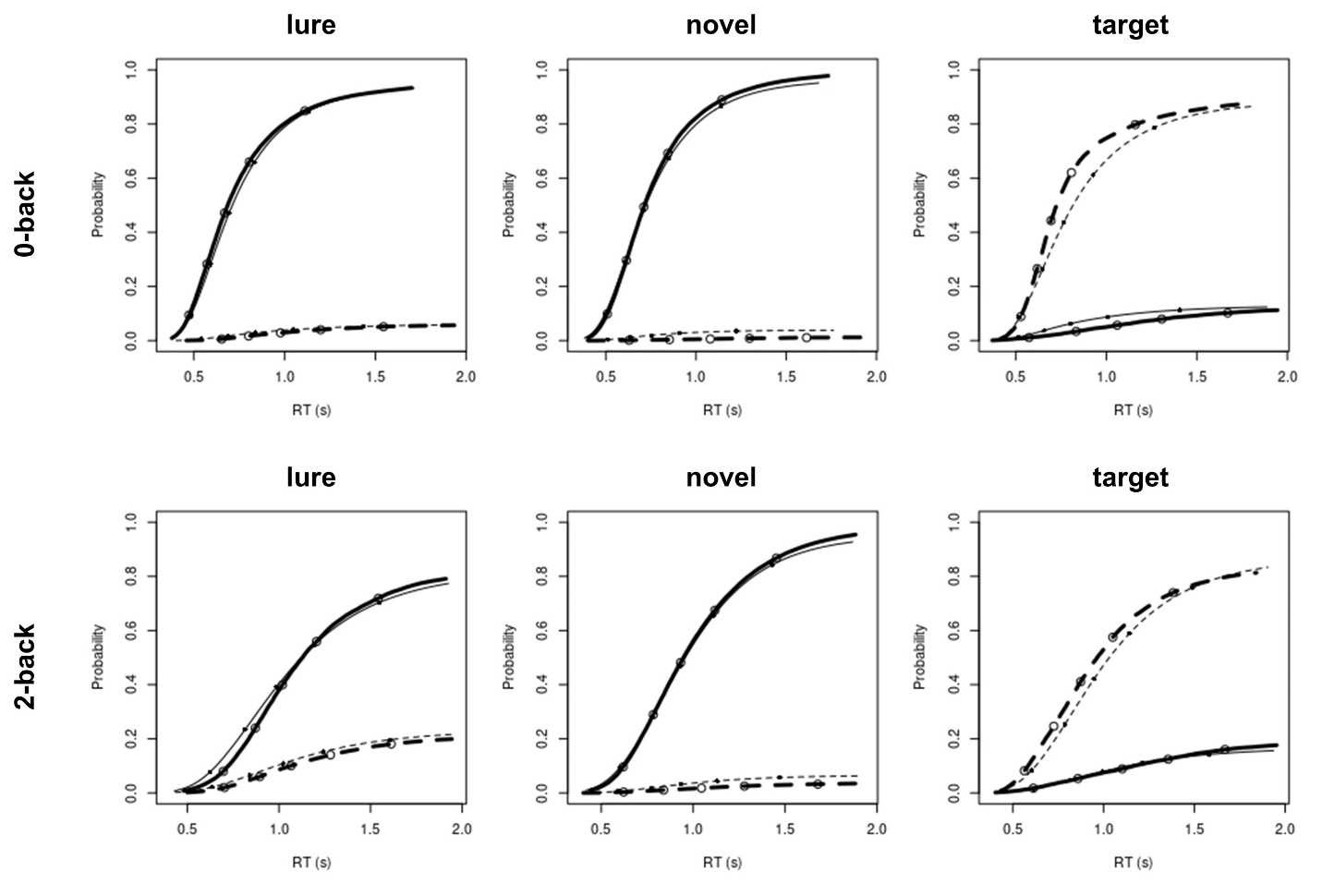

**Supplemental Figure 2.** Posterior predictive plots for diffusion decision model (DDM) fits to the Adolescent Brain Cognitive Development Study (ABCD) n-back task data in each cognitive load and trial condition. Plots display the cumulative probability of a “non-target” (solid line) and “target” (dotted line) response for empirical (thick line) and model-predicted (thin line) data. Specific response time (RT) quantiles (.1, .3, .5, .7, .9) are also displayed for empirical (open dots) and model-predicted (solid dots) data.

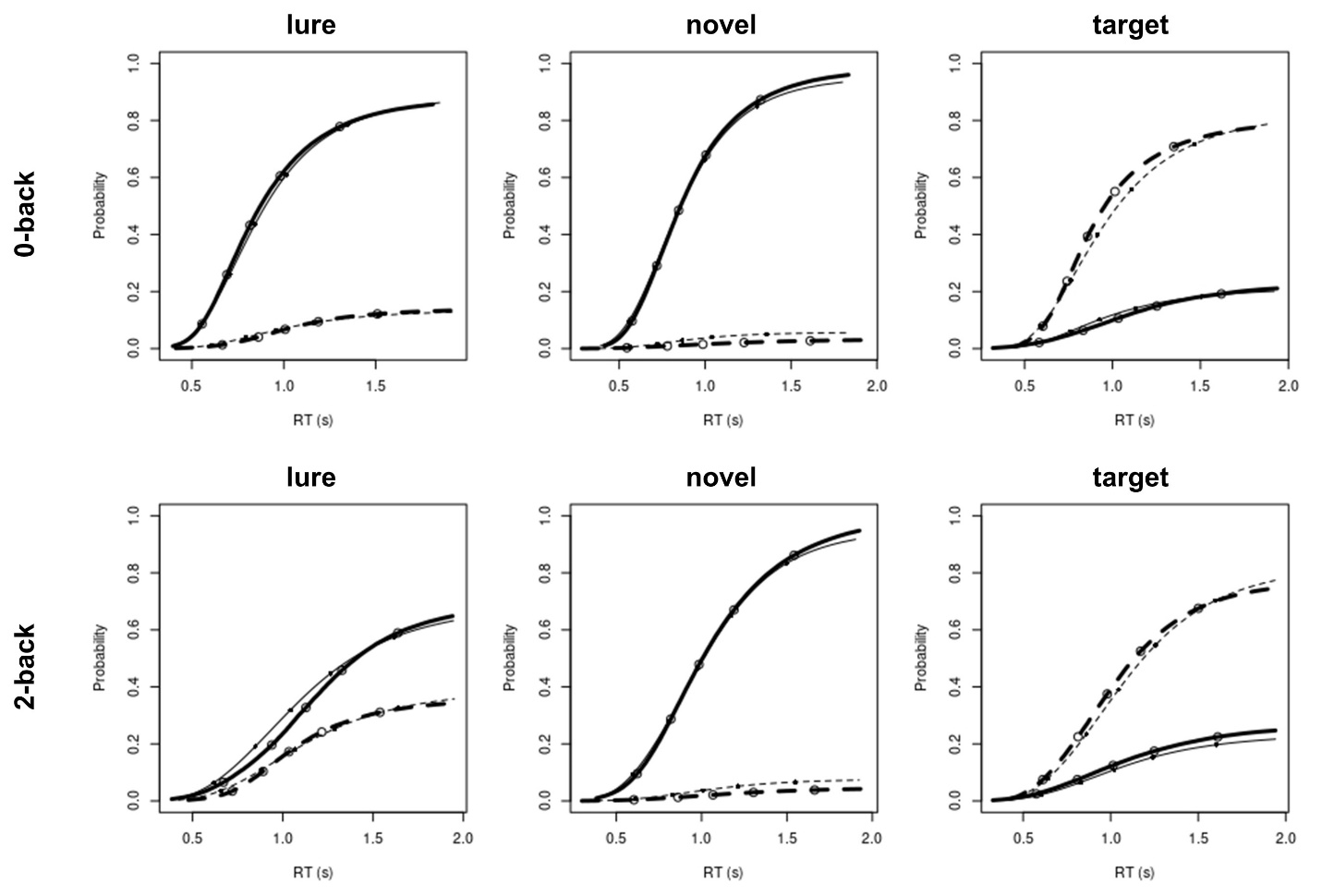

**Supplemental Figure 3.** Visualization of dynamic relations between average task-positive network activation in the 0-back and 2-back conditions and overall efficiency of evidence accumulation (EEA) on the task for the Adolescent Brain Cognitive Development Study (ABCD; left column) and Human Connectome Project (HCP; right column) samples without adjustments for covariates. Values were converted to standardized scores (Z-scores: mean = 0, SD = 1) for interpretability. Individuals’ EEA is represented by the hue of the points, with individuals higher in EEA having darker red hues. Activations of the frontoparietal network (FPN) are shown in the top row while activations of the dorsal attention network (DAN) are shown in the bottom row. Black dotted lines represent the regression line for relations between 0-back and 2-back task activations. Combined with the gray dotted lines representing the average 0-back activation level, the regression lines form four quadrants that denote whether individuals have higher or lower 2-back activation than would be expected given their level of 0-back activation. Bold numbers reflect the average EEA of individuals in each quadrant.

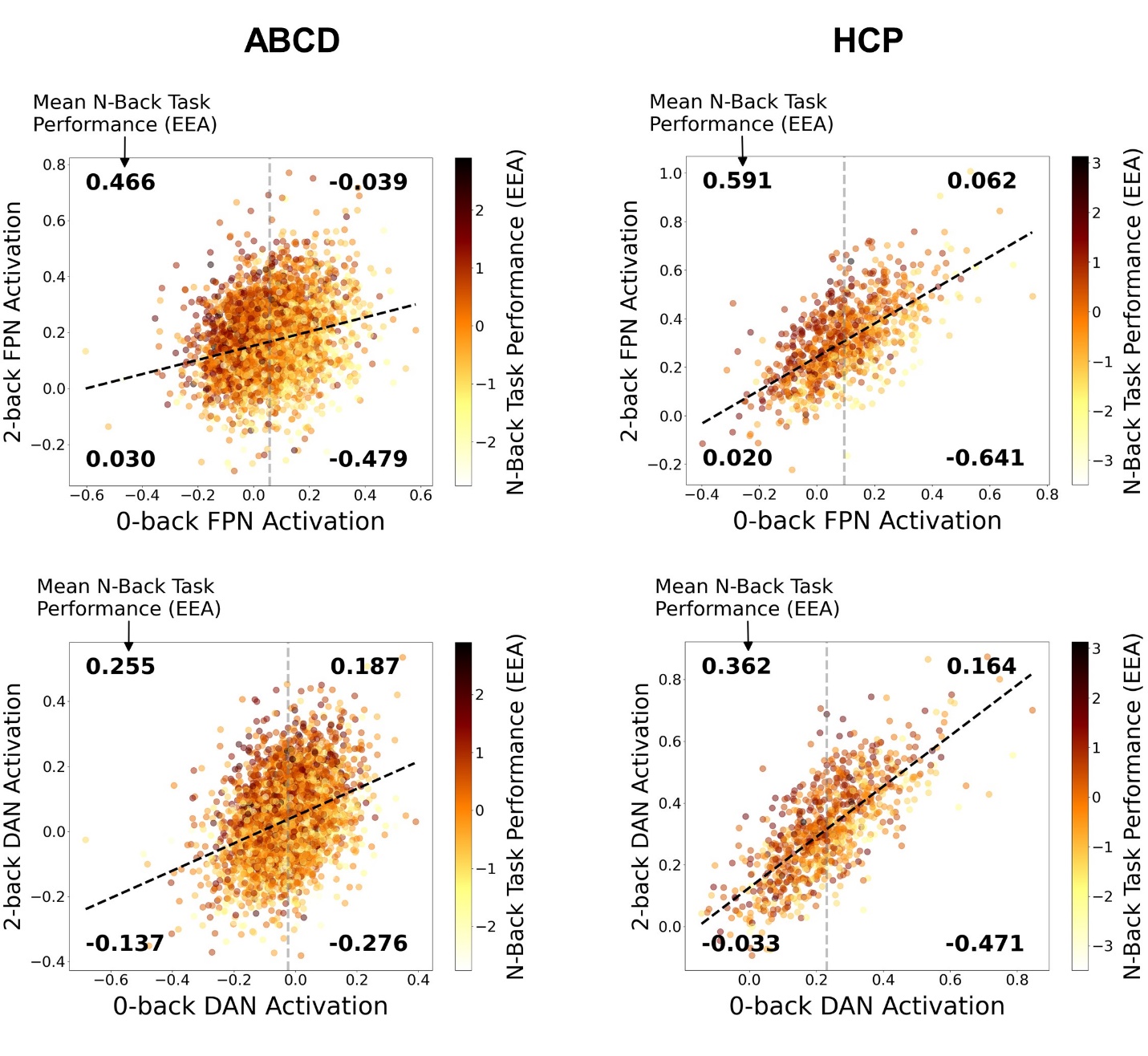

**Supplemental Table 1.** Adolescent Brain Cognitive Development Study (ABCD) and Human Connectome Project (HCP) correlations between efficiency of evidence accumulation (EEA) and average measures of frontoparietal network (FPN) and dorsal attention network (DAN) activation for the 0-back, 2-back and cognitive load (2-0) contrast without adjustments for covariates. 95% confidence intervals, displayed in italics next to each correlation, were estimated using a clustered bootstrapping procedure that accounted for nesting by family and study site.

|  | **ABCD**  ***r*** | **ABCD 95% CI** | | **HCP**  ***r*** | **HCP**  **95% CI** | |
| --- | --- | --- | --- | --- | --- | --- |
| FPN 0 | -.29 | *-.27* | *-.31* | -.33 | *-.27* | *-.39* |
| FPN 2 | .19 | *.22* | *.15* | .09 | *.16* | *.02* |
| FPN 2-0 | .39 | *.42* | *.36* | .50 | *.55* | *.45* |
| DAN 0 | -.06 | *-.03* | *-.09* | -.15 | *-.08* | *-.22* |
| DAN 2 | .21 | *.25* | *.18* | .10 | *.16* | *.03* |
| DAN 2-0 | .25 | *.28* | *.23* | .35 | *.40* | *.29* |
